# Antisense transcription and its roles in adaption to environmental stress in *E. coli*

**DOI:** 10.1101/2023.03.23.533988

**Authors:** Lei Zhao, Ehsan Tabari, Hua Rong, Xia Dong, Di Xue, Zhengchang Su

## Abstract

It has been reported that a highly varying proportion (1% ∼ 93%) of genes in various prokaryotes have antisense RNA (asRNA) transcription. However, the extent of the pervasiveness of asRNA transcription in the well-studied *E. coli* K12 strain has thus far been an issue of debate. Furthermore, very little is known about the expression patterns and functions of asRNAs under various conditions. To fill these gaps, we determined the transcriptomes and proteomes of *E. coli* K12 at multiple time points in five culture conditions using strand-specific RNA-seq, differential RNA-seq, and quantitative mass spectrometry methods. To reduce artifacts of possible transcriptional noise, we identified asRNA using stringent criteria with biological replicate verification and transcription start sites (TSSs) information included. We identified a total of 660 asRNAs, which were generally short and largely condition-dependently transcribed. We found that the proportions of the genes which had asRNA transcription highly depended on the culture conditions and time points. We classified the transcriptional activities of the genes in six transcriptional modes according to their relative levels of asRNA to mRNA. Many genes changed their transcriptional modes at different time points of the culture conditions, and such transitions can be described in a well-defined manner. Intriguingly, the protein levels and mRNA levels of genes in the sense-only/sense-dominant mode were moderately correlated, but the same was not true for genes in the balanced/antisense-dominant mode, in which asRNAs were at a comparable or higher level to mRNAs. These observations were further validated by western blot on candidate genes, where an increase in asRNA transcription diminished gene expression in one case and enhanced it in another. These results suggest that asRNAs may directly or indirectly regulate translation by forming duplexes with cognate mRNAs. Thus, asRNAs may play an important role in the bacterium’s responses to environmental changes during growth and adaption to different environments.

**IMPORTANCE:** The *cis*-antisense RNA (asRNA) is a type of understudied RNA molecules in prokaryotes, which is believed to be important in regulating gene expression. Our current understanding of asRNA is constrained by inconsistent reports about its identification and properties. These discrepancies are partially caused by a lack of sufficient samples, biological replicates, and culture conditions. This study aimed to overcome these disadvantages and identified 660 putative asRNAs using integrated information from strand-specific RNA-seq, differential RNA-seq, and mass spectrometry methods. In addition, we explored the relative expression between asRNAs and sense RNAs and investigated asRNA regulated transcriptional activity changes over different culture conditions and time points. Our work strongly suggests that asRNAs may play a crucial role in bacterium’s responses to environmental changes during growth and adaption to different environments.

## INTRODUCTION

With the advent of high-throughput RNA sequencing (RNA-seq) technologies, our views on the repository of transcriptional molecules have been widely expanded in both eukaryotes and prokaryotes (1, 2). Even the prokaryotic transcriptome, once considered to be relatively simple, turns out to be much more complex than expected (2–5). Besides the traditionally known ribosomal RNAs (rRNAs), messenger RNAs (mRNAs), and transfer RNAs (tRNAs), pervasive noncoding RNAs, and 5’- and 3’-untranslated regions (UTRs) were discovered in a large collection of bacterial species (6). Of these newly uncovered noncoding RNA molecules, *cis*-acting antisense RNAs (asRNAs), encoded on the opposite strand of the coding region of a gene, have been caught in the spotlight as they are complementary to the gene and thus are likely to form duplexes with the cognate mRNA, thereby affecting the mRNA’s function. Many studies found that asRNAs were widespread in taxonomically diverse bacterial lineages, including Gram-positive bacteria such as *Bacillus anthracis* (7), *Bacillus subtilis* (6, 8)*, Listeria monocytogenes* (9), *Staphylococcus aureus* (6), etc. and Gram-negative bacteria, *Escherichia coli* (10–15)*, Salmonella enterica* (12), *Mycoplasma pneumoniae* (8), *Helicobacter pylori* (16), *Pseudomonas aeruginosa* (17), and *Pseudomonas syringae* (18). Such pervasive presence of asRNAs strongly stimulated scientific interest in understanding asRNAs with the expectation of elucidating many aspects of important regulatory functions of asRNAs (19–22).

The earliest asRNA characterization can be traced back to the 1980s when several asRNAs were discovered to regulate plasmid ColE1 (23), plasmid R1 (24), phages (25), and transposons (26). The exploration of asRNA functions was limited at that time, as asRNA regulation was viewed as a rare biological event. In the past 20 years, although the wide adoption of microarray and RNA-seq technologies propelled transcriptome studies, the illustration of asRNA functions is not exploding with accumulating transcriptomic data. Instead, the functions of asRNAs are still unclear-even the authenticity of asRNA became a debate in the scientific community (8). Interestingly, even for one of the most studied model organisms, *E. coli*, reports about asRNA were quite inconsistent. The numbers of asRNAs reported in these studies are in the range of hundreds to thousands. For example, Lybecker et al. studied the double-stranded transcriptome of *E. coli* through an immunoprecipitation technique and identified 316 asRNAs that were produced via RNase III cleaving mRNA-asRNA complex (10). Thomason et al. explored the differential RNA-seq (dRNA-seq) technique on *E. coli* and reported 5,495 antisense transcription start sites (TSSs), though full-length asRNAs were not the focus of the study (26). Dornenburg et al. demonstrated the existence of ∼1,000 asRNAs (11). The inconsistent results cast doubts about the authenticity of asRNA in the model bacterium.

The discrepant results can be partially explained by several factors. Firstly, while the research teams tried to investigate different aspects of asRNAs, they designed the experiments for different purposes, thus the experiment procedures, sequencing library preparation, sequencing platforms, and data analysis methods could all be the sources of bias. Secondly, the definition of an asRNA could also be different from group to group. With varying levels of stringency, an asRNA predicted by one group could fail to meet the standard set by another group. For example, in Dornenburg et al. ’s work, 141 of 1,005 asRNAs overlapping gene bodies were only covered by a single RNA-seq read, thus might be considered as noise transcripts under more stringent criteria (11). Finally, the pipeline and programs used in the analysis could also cause bias. All these factors collectively contribute to the inconsistent results reported.

One could argue that these factors can be hard to avoid, especially for the sequencing-based analysis. However, besides these causes, we also noticed some common issues while we surveyed these asRNA research. First, most studies were based on a limited number of samples. Second, most reports lacked the verification from biological replicates, thus downgrading the confidence in the conclusions. Third, most bacterial cells were cultured in and collected from one or two mediums, mainly the rich medium providing sufficient nutrients. The lack of diverse culture conditions may lead to an underestimation of asRNAs, as it has been reported that asRNAs may show different expression patterns under different environments (6, 27).

To overcome these shortcomings, in this study, we conducted our research with a systems approach. We collected *E. coli* cells from five different culture conditions across 22 growth time points and prepared directional RNA-seq libraries with a modified dRNA-seq method (16, 28). dRNA-seq can help determine TSSs, thereby infusing another layer of confidence in asRNA predictions. We also prepared at least two biological replicates for each growth time point. With the profiled 660 asRNAs, we found that asRNA transcription in *E. coli* is pervasive yet highly variable and dynamic, that asRNA expression displayed a culture-dependent manner, and that many genes changed their relative levels of asRNAs to the mRNAs at different time points under different conditions, suggesting asRNAs might play crucial roles in the bacterium’s responses to environmental changes.

## RESULTS

### Our RNA-seq and dRNA-seq libraries are sufficient and of high quality

To overcome the intrinsic difficulty to study asRNAs with traditional disruptive methods that would affect the sense RNA transcription unavoidably, we designed our study using a systems approach. Using a combination of strand-specific RNA-seq, dRNA-seq, and mass spectrometry methods, we profiled the transcriptome and proteome of *E. coli* K-12 cells at different time points under five different culture conditions, including rich medium Luria Broth (LB), minimum nutrient medium (MOPS), heat shock (HS) in MOPS, carbon starvation (MC), and phosphorus starvation (MP). For the LB samples, we collected the bacteria when OD_600_ reached 0.5, 1.0, 2.0, 3.0, and 4.0, respectively, representing the early (OD_600_ = 0.5), middle (OD_600_ = 1.0 and 2.0), and log phases (OD_600_ = 3.0 and 4.0) of cell growth in the medium. For the other four culture conditions, we started culturing the cells in LB and transferred them to one of the four other stress conditions when OD_600_ = 1.0. Then, we collected the cells at different time points from each of these stress conditions.

In total, we generated 110 strand-specific RNA-seq libraries. Among them, 88 were prepared with a modified dRNA-seq method that helps enrich the 5’-ends, and thus the TSSs of the transcripts. Generally, extracted RNAs were equally split into two parts with one half having Terminator™ 5′-Phosphate-Dependent Exonuclease (TEX) added and the other half without. TEX is a processive 5′-end to 3′-end exonuclease that digests processed RNAs having a 5′ monophosphate group, but not the unprocessed RNAs having a 5′-triphosphate, 5′-cap, or 5′-hydroxyl group, so the unprocessed transcripts can be enriched to magnify the TSS information. We generated a total of 88 libraries for two biological replicates for each type (+TEX and -TEX) of the libraries at each time point (2 replicates x 2 types of libraries x 22 time points). In addition, we also included in our analyses 22 RNA-seq libraries generated from rRNA-depleted samples collected in the same culture conditions (13), which can be viewed as -TEX libraries in the analysis. A full list of conditions, time points, and the sample size was summarized in Table 1.

**Table 1.**
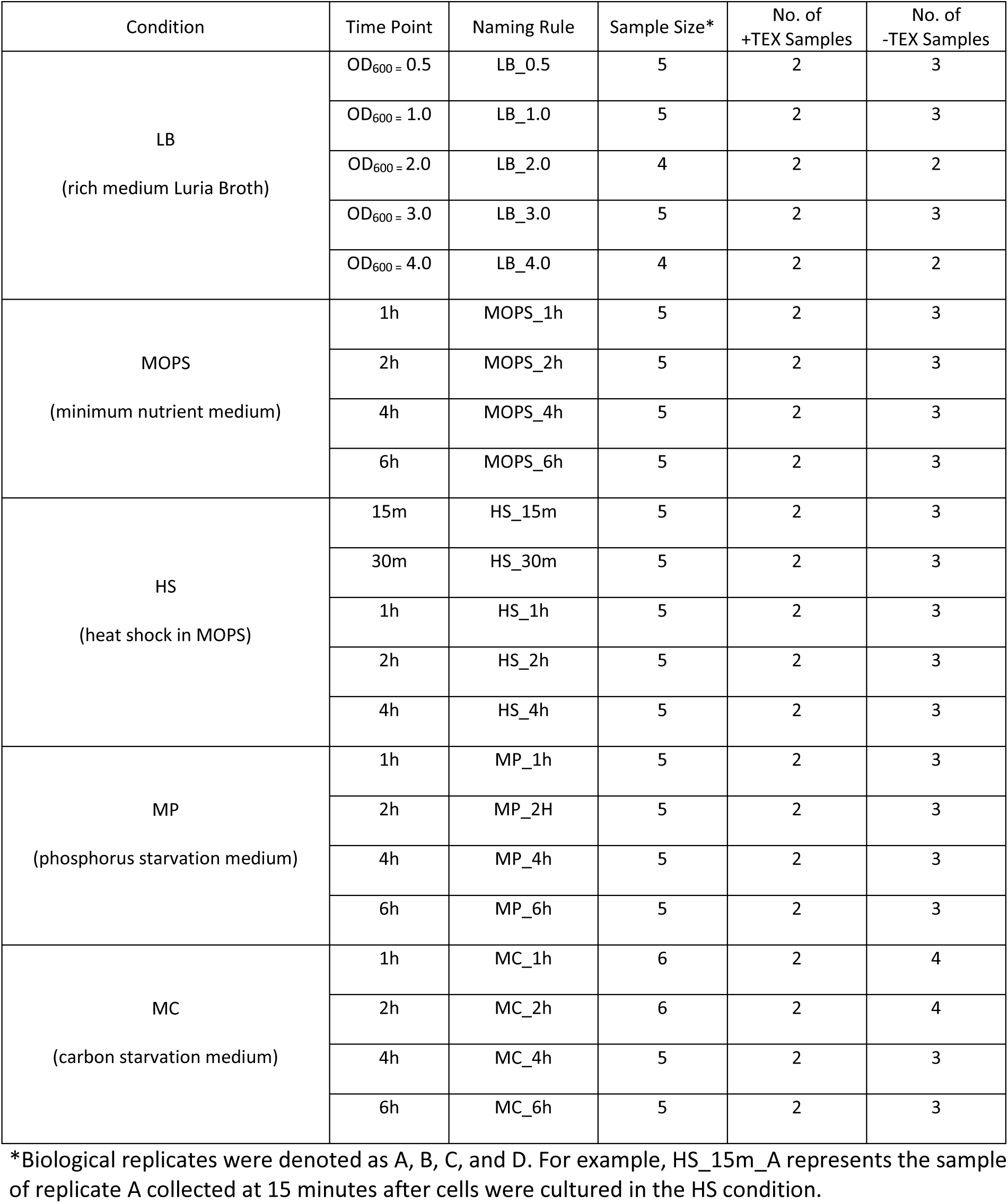
Summary of culture conditions, sample collection times, and sample sizes

| Condition | Time Point | Naming Rule | Sample Size* | No. of +TEX Samples | No. of -TEX Samples |
| --- | --- | --- | --- | --- | --- |
| LB<br>(rich medium Luria Broth) | OD<sub>600</sub> = 0.5 | LB_0.5 | 5 | 2 | 3 |
|  | OD<sub>600</sub> = 1.0 | LB_1.0 | 5 | 2 | 3 |
|  | OD<sub>600</sub> = 2.0 | LB_2.0 | 4 | 2 | 2 |
|  | OD<sub>600</sub> = 3.0 | LB_3.0 | 5 | 2 | 3 |
|  | OD<sub>600</sub> = 4.0 | LB_4.0 | 4 | 2 | 2 |
| MOPS<br>(minimum nutrient medium) | 1h | MOPS_1h | 5 | 2 | 3 |
|  | 2h | MOPS_2h | 5 | 2 | 3 |
|  | 4h | MOPS_4h | 5 | 2 | 3 |
|  | 6h | MOPS_6h | 5 | 2 | 3 |
| HS<br>(heat shock in MOPS) | 15m | HS_15m | 5 | 2 | 3 |
|  | 30m | HS_30m | 5 | 2 | 3 |
|  | 1h | HS_1h | 5 | 2 | 3 |
|  | 2h | HS_2h | 5 | 2 | 3 |
|  | 4h | HS_4h | 5 | 2 | 3 |
| MP<br>(phosphorus starvation medium) | 1h | MP_1h | 5 | 2 | 3 |
|  | 2h | MP_2H | 5 | 2 | 3 |
|  | 4h | MP_4h | 5 | 2 | 3 |
|  | 6h | MP_6h | 5 | 2 | 3 |
| MC<br>(carbon starvation medium) | 1h | MC_1h | 6 | 2 | 4 |
|  | 2h | MC_2h | 6 | 2 | 4 |
|  | 4h | MC_4h | 5 | 2 | 3 |
|  | 6h | MC_6h | 5 | 2 | 3 |

\*Biological replicates were denoted as A, B, C, and D. For example, HS\_15m\_A represents the sample of replicate A collected at 15 minutes after cells were cultured in the HS condition.

After quality control and 3’-adapter clipping, we collected a total of about 1.16 billion reads from these 110 libraries with 44 of them being the 5’-enriched libraries (+TEX) and the remaining 66 being the rRNA-depleted libraries (-TEX). The vast majority (74.0%∼98.8%) of the reads of the libraries except one (-TEX MC_6h_B, 50.5%) can be mapped to the *E. coli* K-12 genome, suggesting that our experiments yielded high-quality data. We further processed the aligned reads by preserving only the unambiguously mapped ones. The number of uniquely mapped reads was between 730, 371 (-TEX LB_0.5_A) and 10,133,642 (-TEX MP_4h_C) and the percentages of these reads were between 4.7% (-TEX MC_6h_C) and 38.2% (-TEX MP_4h_C) with a median of 16.9%. The highly varying unique mapping rates are due to the fact that most of the total RNAs are rRNAs, and our rRNAs depletion kits have varying efficiency. However, the number of uniquely mapped reads (Table S1) is sufficient for our analyses as indicated as follows.

### We found a total of 660 antisense RNAs using stringent criteria

In our libraries, the average percentage of antisense reads is ∼ 1.1%. For most genes, the asRNA expression was lower than its counterpart sense RNA expression, so the number of antisense sequencing reads is fewer than the number of sense sequencing reads, which is consistent with previous studies (29, 30). With these antisense reads, we assembled a total of 660 antisense RNAs using our stringent criteria, associated with 816 genes (19.9% of the *E. coli* genes) (Table S2). Most (77.4%) of these asRNAs overlapped with one gene while the remaining 22.6% overlapped with multiple genes (Fig.1A). 17.7% of asRNAs were identified at certain time points only, while the remaining 82.3% were observed in at least two time points (Fig.1B). In consistent with earlier reports that asRNA lengths varied from ten to thousands of nucleotides (10), lengths of our predicted asRNAs range from 14 to 5,332 nucleotides (nt). Lengths of asRNAs and those of the cognate genes were moderately correlated, as indicated by Spearman correlation coefficient ρ value of 0.45 (Fig.1C), and asRNAs were generally shorter than their cognate genes (Fig.1D).

**Fig.1.**
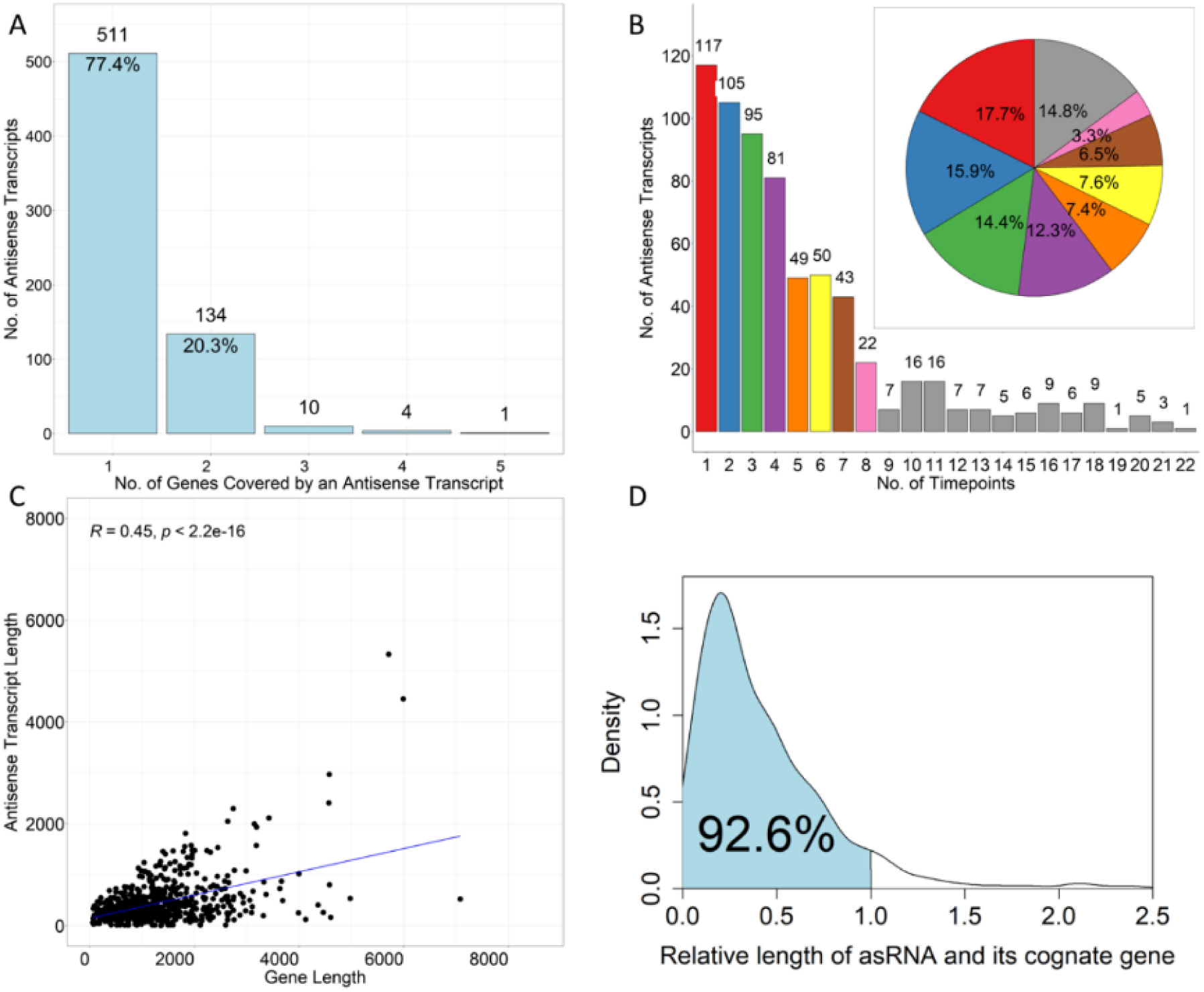
Properties of the predicted antisense RNAs. (A) Counts of antisense RNAs that overlapped with different numbers of genes. (B) The number of time points in which the same antisense transcripts were observed. The inset pie chart shows the percent of the number of antisense transcripts observed in different numbers of time points. (C) The relationship between the lengths of antisense transcripts and those of the cognate genes. (D) The density of the ratios of the lengths of asRNAs over their cognate genes.

### asRNAs are more likely to initiate from within the ORF than the UTRs of a gene

Earlier studies showed that an asRNA can overlap part of a gene, an entire gene, or even several genes and that the TSS of the asRNA can be located within the Open Reading Frame (ORF) or the UTRs of the gene (20, 22, 31). With the overlapping and contiguous genes in the compact prokaryotic genomes, the relative locations of asRNAs to their cognate genes can be complex, as summarized by Lybecker et al. (10) that there were at least 10 scenarios for asRNAs’ locations relative to the annotated ORFs. However, in at least two of these scenarios, the identified asRNAs may not be bona fide, as the mRNA for one gene might be an asRNA for its convergent or divergent neighbor gene. Thus, the “excludon” concept was proposed to describe this situation (32, 33). Our predicted asRNAs’ relative locations fit into the remaining eight scenarios, with full-ORF being the most frequent when an asRNA overlaps at least half of the cognate sense transcript (Fig.2A).

**Fig.2.**
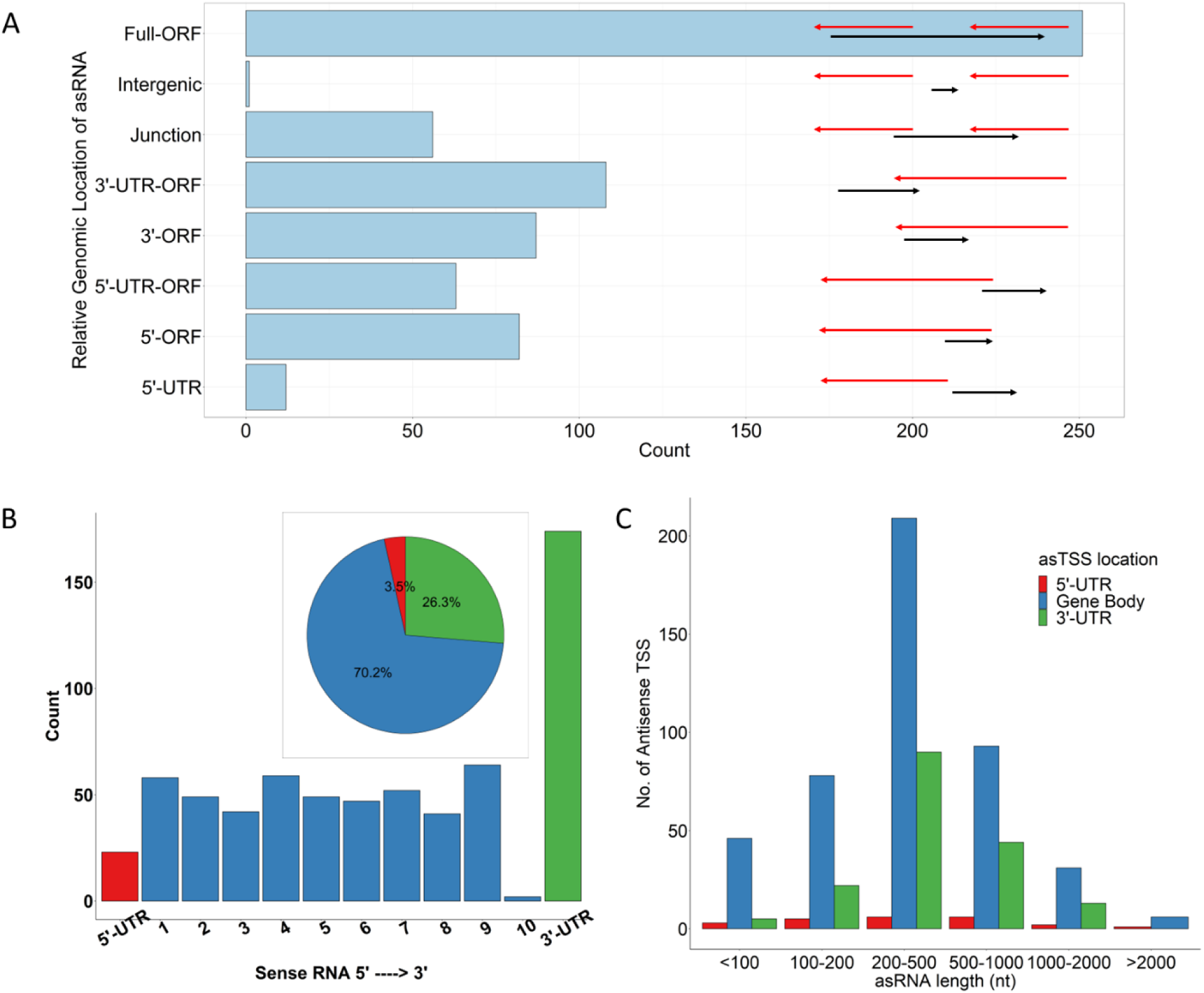
Genomic locations of the predicted asRNAs. (A) Schematic illustration of the eight categories of asRNAs based on their relative locations to the ORFs and the counts in each category. The schemes were modified from Lybecker et al.’s work (10) with “divergent” and “convergent” categories excluded. Red arrows represent the genes and their orientations and so do the black arrows for the asRNAs. (B) Counts of asTSSs in the 5’-UTR, gene body, and 3’-UTR. The numbers from 1 to 10 on the horizontal axis are the equally divided bins for each gene body. The inset shows the percentage of the asRNAs with their TSS falling in the 5’-UTR, gene body, and 3’-UTR. (C) Counts of asRNAs with a length in the indicated regions having their TSS falling in the 5’-UTR, gene body, and 3’-UTR.

In our analysis, we use antisense TSSs as one of the major pieces of evidence for asRNA identification. The TSSs were identified based on the genome-wide comparison at single nucleotide resolution between the +TEX and -TEX libraries generated by the dRNA-seq techniques. The antisense TSSs were determined by the TSSPredator program (34) and supplemented by upshift peaks between consecutive nucleotides in the genome with manual review. For our 660 predicted asRNAs, 70.0% were initiated from within ORFs, while 26.3%, 3.5%, and 0.2% were from within the 3’-UTRs, 5’-UTRs, and intergenic regions, respectively (Fig.2B), suggesting that asRNAs preferred to initiate from within the ORF than the UTRs of a gene. We then equally divided the gene body into 10 bins and calculated the frequency of asTSSs in each bin. The count shows that there is no loci preference for asTSSs within the gene except for the 10^th^ bin at the 3’-end having a low count, indicating that when an asRNA initializes around the 3’-end of a gene, it tends to use the 3’UTR rather than the gene body at the 3’-end (Fig.2B). To see whether or not the length of an asRNA is related to the location of its TSS in the gene, we divided asRNAs into several groups by their lengths. As shown in Fig.2C, asRNAs prefer to initialize from within the ORF or UTRs, regardless of their length. Thus, asRNA lengths have little to do with the asRNAs’ initialization preference.

### Antisense RNAs are expressed in a culture condition-dependent manner

Next, we explored the observation that the predicted asRNAs were expressed in different culture conditions. As shown in Fig.3A, although 60 (9.1%) asRNAs were observed in all five culture conditions, the remaining was only expressed in a certain number of conditions with 276 (41.8%) asRNAs in only one condition. Interestingly, most of these uniquely expressed asRNAs occurred in HS and MP conditions (Fig.3B). These results indicate that the expression of most asRNAs was highly condition-dependent, thus they might play a role in the adaption of the bacteria to the respective culture conditions.

**Fig.3.**
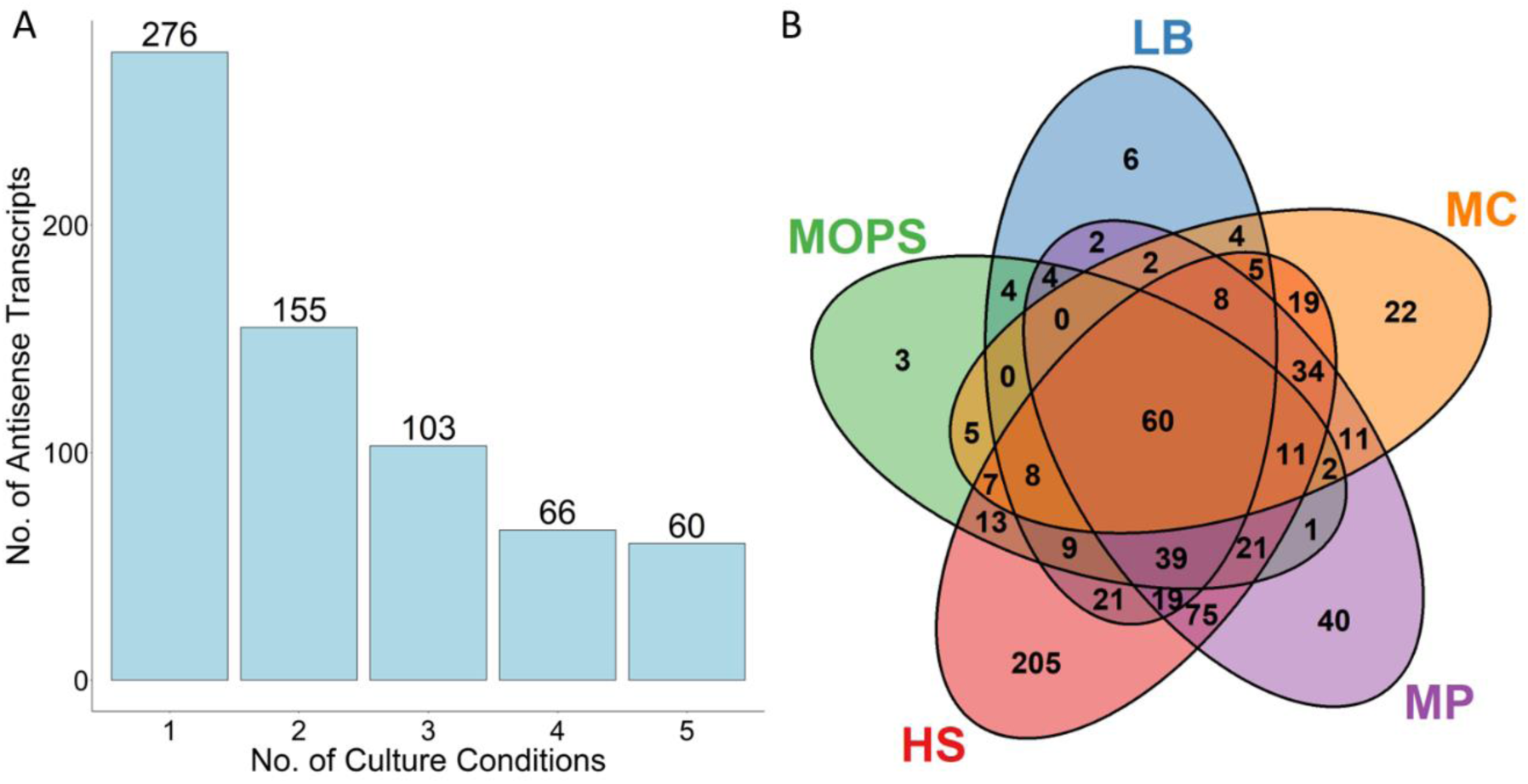
Culture-dependent expression of the predicted asRNAs. (A) The number of asRNAs expressed in the number of culture conditions on the horizontal axis. (B) Venn diagram of asRNAs in the five culture conditions showing shared and unique asRNAs.

### Transcriptional events at a locus can be categorized into six modes

The known molecular mechanisms of asRNA regulation can be classified into four types including transcription inference, attenuation, stability alteration, and translation inhibition (6, 20, 32). Except for inference, all other three mechanisms require physical RNA-RNA interactions, therefore, it is widely believed that asRNAs execute their functions by forming RNA duplex with the sense RNAs counterparts (35, 36). To better understand the relationship between sense RNA and asRNAs, it is necessary to quantify them in the first place. Here, we proposed a parameter, ɣ, define as the ratio of the expression level of an asRNA and that of its counterpart mRNA at the logarithm scale, to explore their relationships. Based on the ɣ value, we categorized transcription events at a gene locus in six modes as defined in Table 2, namely, sense-only, sense-dominant, balanced, antisense-dominant, antisense-only, and silent. As shown in Fig.4, the distribution of ɣ values displays a strongly left-skewed, bell-shaped curve, indicating that most gene loci are in sense-only and sense-dominant modes, i.e., the asRNA expression level at a gene locus is generally lower than the mRNA expression level, which is consistent with earlier reports (29, 30). However, a considerable proportion of gene loci are in balanced, antisense-dominant, and antisense-only modes. This pattern of distribution of ɣ values is also seen for gene loci at all the time points of all the five culture conditions, although the time-course-dependent shift of the distributions also is clear (Fig.4), suggesting that gene loci might change their transcriptional modes over time.

**Table 2.** Definitions of the transcriptional modes at a gene locus

| Transcriptional Mode | Acronyms | Defining Condition $\gamma = \log(\text{asRNA}/\text{mRNA})$ |
| --- | --- | --- |
| Sense-dominant | SD | $\gamma < -0.5$ and Expression of asRNA $\neq 0$ |
| Sense-only | SN | $\gamma < -0.5$ and Expression of asRNA $= 0$ |
| Antisense-dominant | AD | $\gamma > 0.5$ and Expression of sense RNA $\neq 0$ |
| Antisense-only | AN | $\gamma > 0.5$ and Expression of sense RNA $= 0$ |
| Balanced | BAL | $-0.5 \leq \gamma \leq 0.5$ |
| Silent | SIL | Expression of sense RNA $= 0$ and<br>Expression of asRNA $= 0$ |

**Fig.4.**
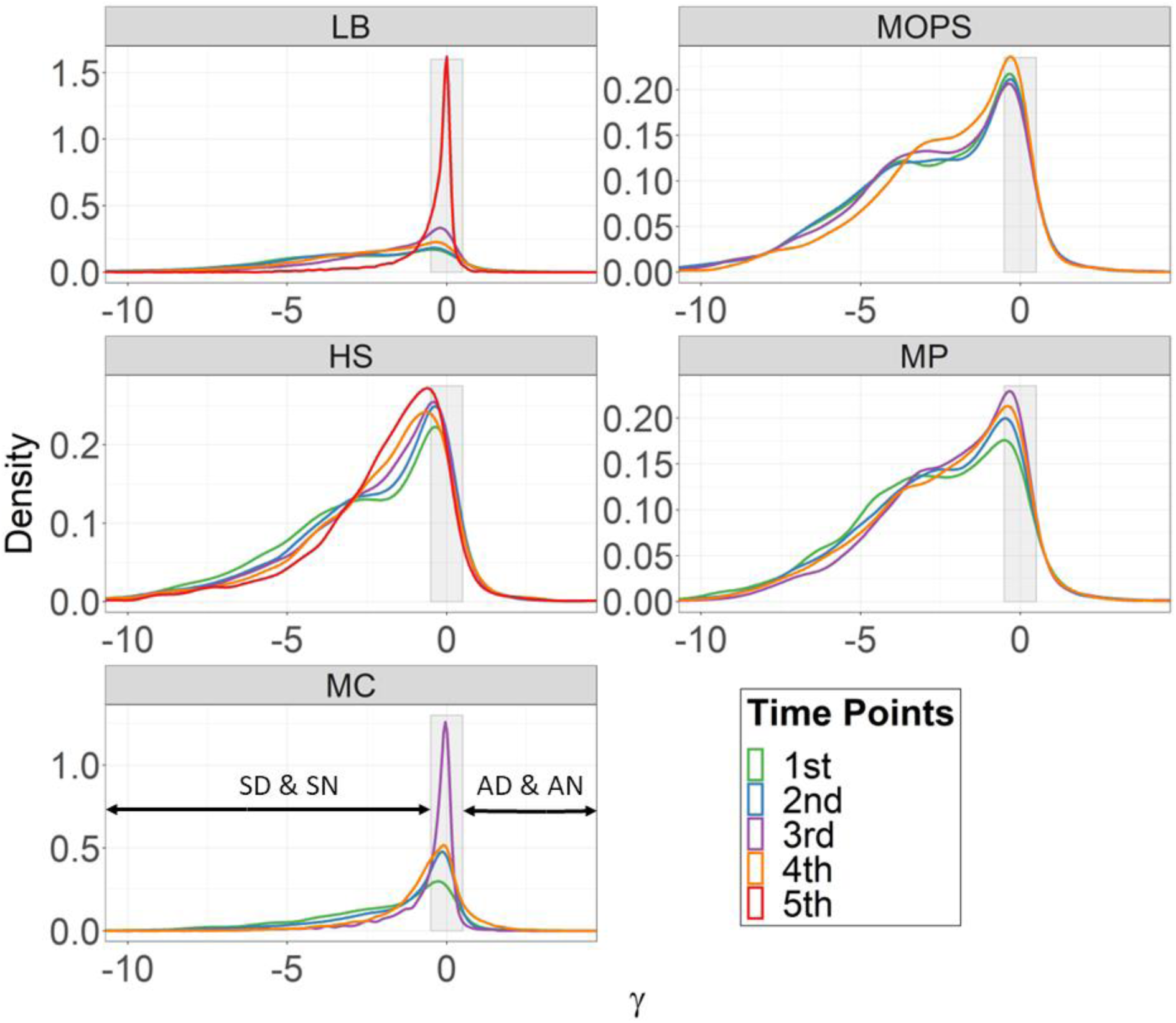
Relative expression levels of asRNAs and their cognate sense RNAs. Density plots of the ɣ values of gene loci at each time point for each culture condition. The gray rectangle between - 0.5 and 0.5 indicates the “balanced” mode whereas asRNA and their cognate sense RNA express similarly, including the silent mode. The lines in each plot are colored by time points.

### Transcriptional mode changes at different conditions/time points

To further investigate the dynamic changes of transcriptional modes of gene loci over time under each culture condition, we calculated the probability that gene loci in a transcriptional mode at a time point would change to a different transcriptional mode or remain in the same transcriptional mode at the next time point. As shown in Fig.5A, although most gene loci remain in the same transcriptional mode between two adjacent time points under all five culture conditions as indicated by the antidiagonal in each heatmap plot, still many gene loci changed their transcriptional modes at certain time points. For example, gene loci in the antisense-dominant mode at LB 4^th^ (LB_3.0) were more likely to change to the balanced or silent modes at LB 5^th^ (LB_4.0) than remaining in the same mode, and gene loci in the sense-only at HS 3^rd^ (HS_30m) were more likely to change to the sense dominant mode at HS 4^th^ (HS_1h) than remaining in the same mode. The number of genes that changed their transcriptional modes was quite different for each culture condition. As shown in Fig.5B, the number of genes changing their transcriptional modes largely increased over time in the LB condition. In MOPS, HS, and MP conditions this number showed similar patterns experiencing a slight decrease at first and then an increase at the end. Interestingly, some gene loci constantly changed their transcriptional modes. For example, in the LB condition, 324 gene loci changed their transcriptional modes between every two adjacent sampling time points. The numbers of gene loci that constantly changed transcriptional modes in the other four conditions were 99 (MOPS), 21 (HS), 103 (MP), and 182 (MC) (Fig.5C). Lastly, we checked the overlap for the gene loci that constantly changed transcriptional modes in all the conditions and found no such gene locus (Fig.5D).

**Fig.5.**
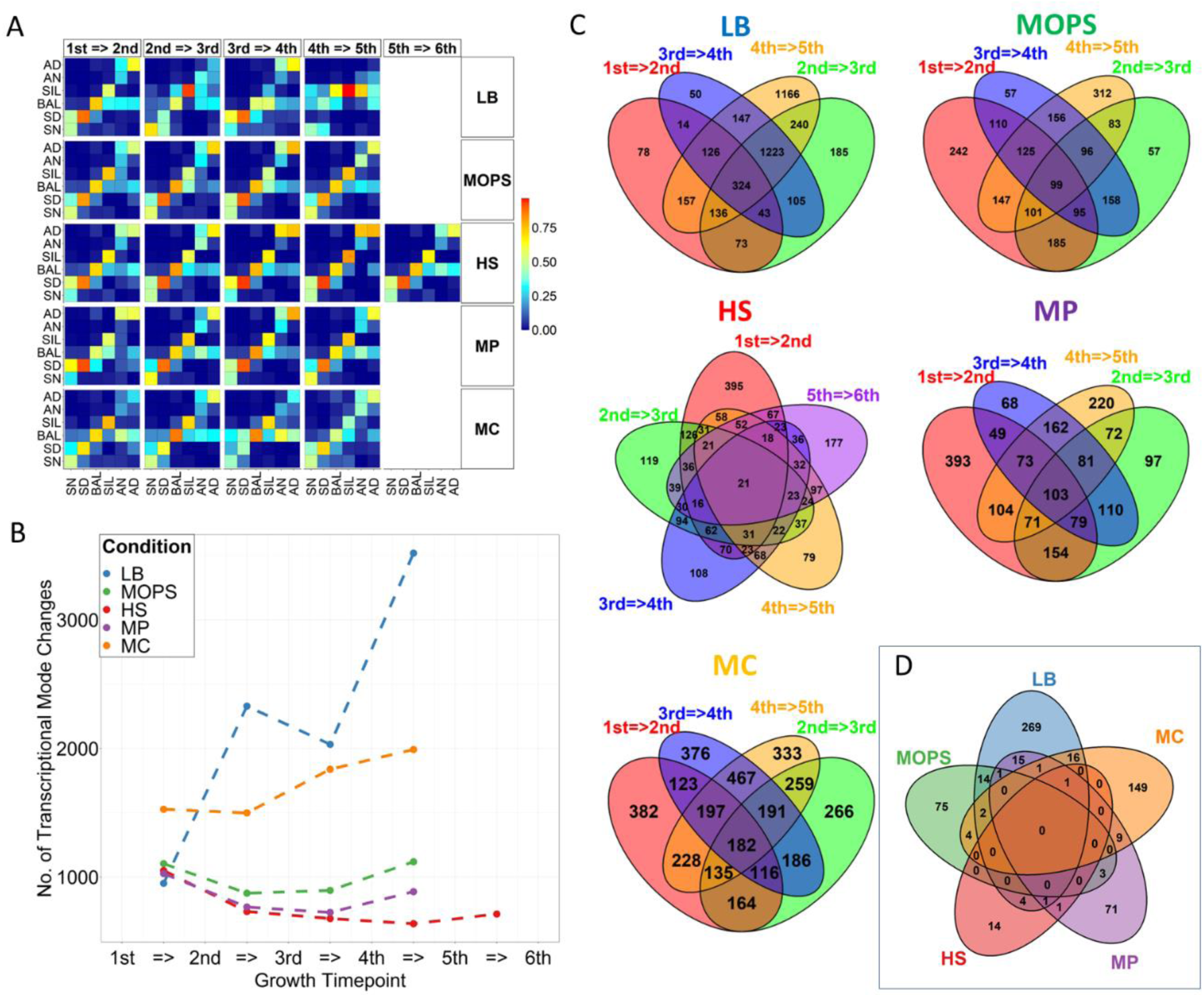
Transitions of transcriptional modes of loci between adjacent sampling time points under each culture condition. (A) Heatmap of the probability that gene loci in a transcriptional mode at a time point would change to a different transcriptional mode or remain in the same transcriptional mode at the next sampling time point (from the x-axis to the y-axis). (B) The number of gene loci that change their transcriptional modes between two adjacent time points. (C) Venn diagram for each condition showing the numbers of shared and unique gene loci having transcriptional mode changes. (D) Venn diagram showing the numbers of shared and unique gene loci having transcriptional mode changes in the five conditions. In plots A, B, and C, the 1^st^ time point in the LB condition refers to LB_0.5, and in the other four conditions refers to LB_1.0, from which the cells were transferred to the indicated culture conditions.

As an example, Fig.6A showed the transcriptional mode changes at different time points in the HS condition for the *sulA* gene encoding the cell division inhibitor. At LB_1.0 (OD_600_=1.0 in LB), the locus is under the balanced transcriptional mode. After transferring to the HS condition, the locus initially (HS_15m, HS_30m) increased mRNA expression levels, thus transitioning to the sense-dominant mode. At HS_1h, its asRNA increased, and it transitioned to the balanced mode. However, at HS_2h, the locus transitioned to antisense-dominant mode and remained in the mode until HS_4h. Another example, as shown in Fig.6B, displayed how the *uspF* gene encoding the universal stress response protein changed its transcriptional modes at different stages in HS. At LB_1.0 (OD_600_=1.0 in LB), the locus was under the sense-dominant mode. After the cells were transferred to the HS condition, it initially (HS_15m) remained in the sense-dominant mode, and then changed to balanced modes thereafter. To see how the protein levels of SulA and UspF change during the time course, we monitored their expression using Western blots. As shown in Fig.6C, the SulA protein that plays a role in SOS responses (37), showed a decrease in the expression in the two later time points under heat shock. While this phenomenon has been noted earlier (38), little is known about the mechanism of the expression reduction. As shown in Fig.6A, there were minimal changes in the mRNA levels of *sulA* at multiple time points under heat shock, however, its antisense expression levels dramatically increased 4 hours after the onset of heat shock, resulting in a transition from the sense-dominant mode at the first two time-points to the antisense-dominant mode at the later time-points. Hence, we hypothesize that the antisense transcription might play an inhibitory role in the expression of the protein. Furthermore, it has been demonstrated that *uspF*, which promotes the adhesion of cells at the expense of motility (39, 40), is upregulated under a glucose-limited condition (41). As shown in Fig.6D, *uspF* was also upregulated under heat shock. Interestingly, the mRNA level of *uspF* remained almost the same from HS_15m to HS_4h, while its antisense level increased, resulting in a transition from the sense-dominant mode in LB_1.0 and HS_15m to the balanced mode in the later stages of heat shock. Thus, this antisense transcript might also enhance the protein expression.

**Fig.6.**
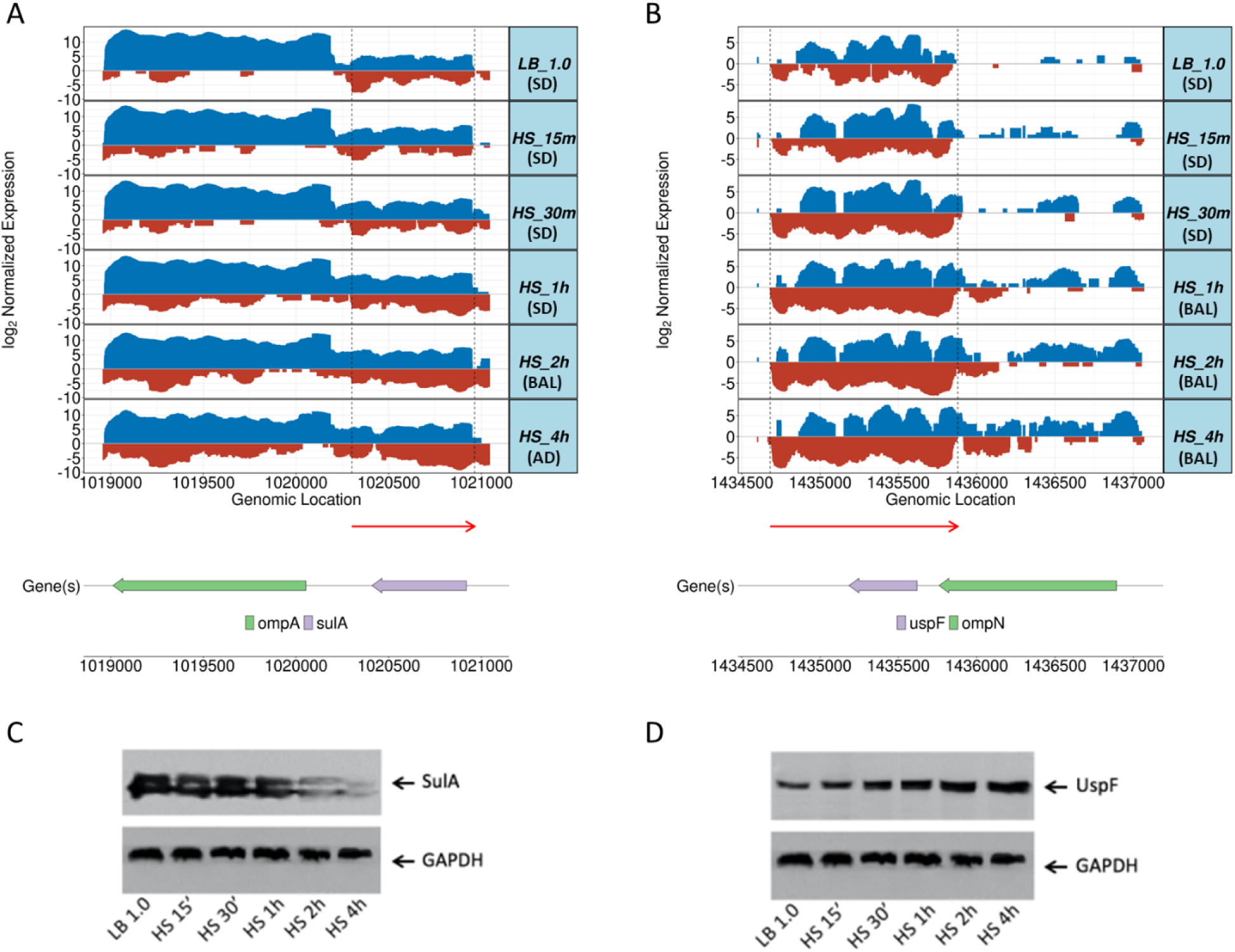
Two examples of transcriptional mode changes under heat shock stress. (A) The *sulA* locus changed its balanced transcriptional mode in LB to the sense-dominant mode, balanced mode, and antisense-dominant mode after being transferred to the heat shock condition. (B) The *uspF* locus changed its sense-dominant transcriptional mode in LB to balanced mode after being transferred to the heat shock condition. In (A) and (B), the sense read coverages are colored in blue with a positive expression value and the antisense reads are colored in red with a negative expression value for display purposes. Identified asRNAs were indicated by the red arrows below the expression plots, with 5’ to 3’ direction. (C) Western blot verification of the expression of the SulA protein under heat shock stress condition. (D) Western blot verification of the expression of the UspF protein under heat shock stress condition. In (C) and (D) GAPDH served as the standard control.

### Protein levels of genes are stoichiometrically affected by the counterpart asRNAs

To see the possible effects of asRNA transcription on the protein expression of the associated genes, we quantified the levels of detected proteins in terms of the number of peptides per hundred amino acids (NPPH, see Materials and Methods). The peptides detected for proteins of the technical replicates for the same samples and biological replicates for the same sampling time points were highly repeatable, so we also pooled the results from the technical and biological replicates for further analysis. We detected 1,270 (in MC_6h) to 1,951 (in LB_1.0) proteins at each time point. Under each culture condition, the protein and mRNA levels were correlated differently for the loci in different transcription modes as indicated by spearman correlation coefficients. As shown in Fig. 7 for the MOPS condition, we observed higher positive correlations for loci in sense-dominant and sense-only transcriptional modes, lower positive or negative correlations for loci in balanced mode and a more complicated scenario for loci in antisense-dominant mode largely due to lack of enough genes in this mode. Similar patterns were also observed in four other conditions (Figure S1). These patterns suggest the increased level of asRNA might disrupt the correlation between mRNA and protein levels in a stoichiometric manner, i.e., depending on the relative abundance of asRNA and mRNA in the bacterial cells.

**Fig.7.**
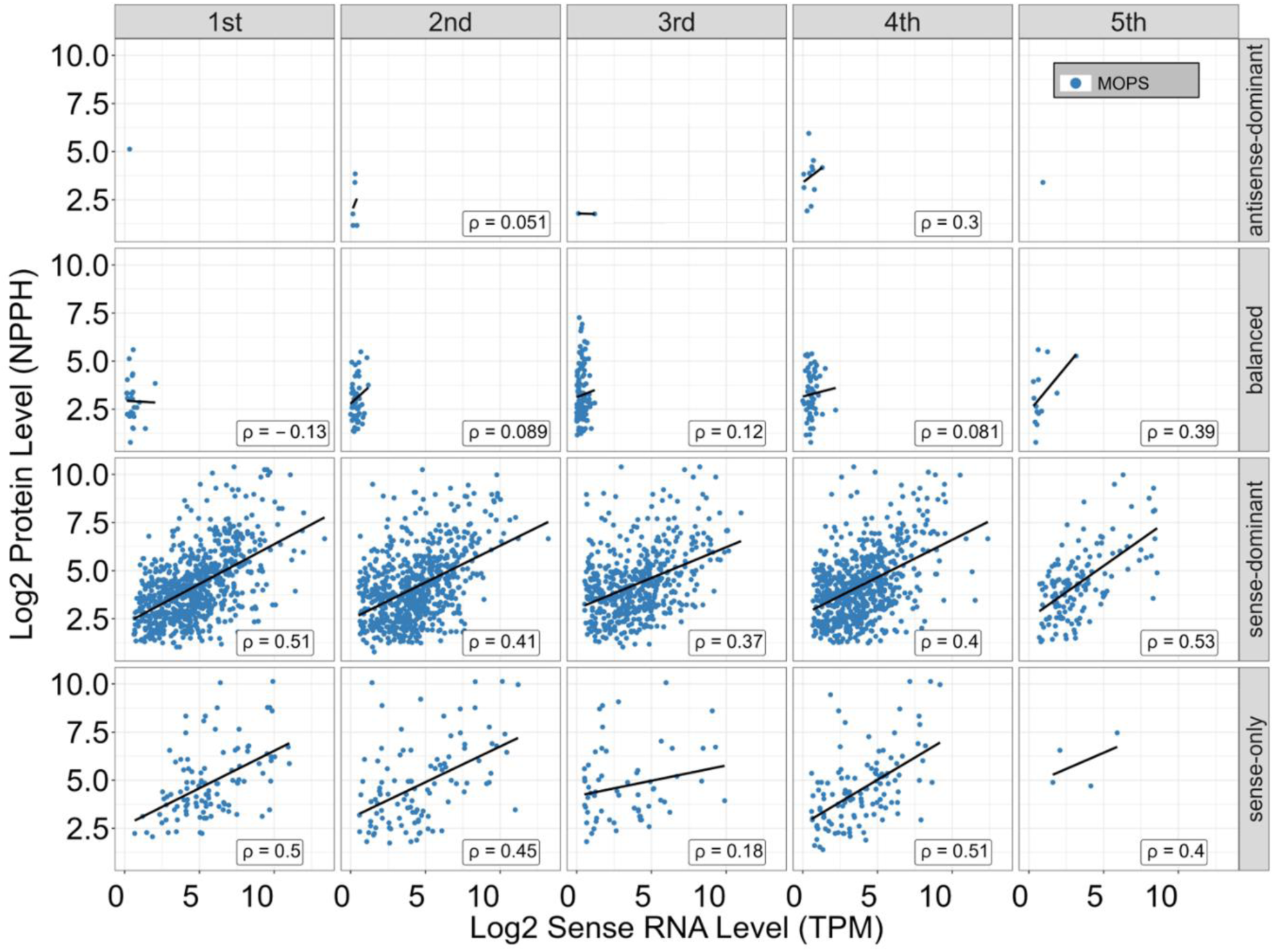
Relationship between the sense transcription levels and protein levels of genes under MOPS culture condition. Correlations between protein levels and mRNA levels of genes in different transcriptional modes (rows) and at different time points (columns). Each data point represents one gene. The Spearman correlation coefficients ρ between the protein levels and mRNA levels of genes are shown in each plot.

## DISCUSSION

With the wide adoption of RNA-seq techniques, mounting evidence suggests pervasive antisense transcriptions in almost all domains of life (1, 2, 22). Specifically for prokaryotes, accumulating reports aimed to address the properties and functions of asRNAs (6, 20, 35, 36). Studies have shown that a highly varying portion of genes in various prokaryotes have asRNA transcription (29), however, it is not clear whether these differences are due to biological or technical variations or both. The reports about the number of asRNAs expressed in a bacterial culture are rather inconsistent. This discrepancy is caused at least partially because different experimental procedures, different sequencing strategies, and different definitions of asRNAs were employed by different groups (10–13, 42). On the other hand, some fixable issues were pervasively existing in these studies. Most of these studies were done based on limited sample size, culture conditions, and often with only one biological replicate (10–12, 14).

In this study, we analyzed the transcriptomes and proteomes in *E. coli* K12 at different growth phases/time points under five culture conditions using a strand-specific RNA-seq method, a dRNA-seq method, and a quantitative mass spectrometry method. To our knowledge, this is the study having the largest sample size so far. The resulting data allowed us to systematically analyze the pervasiveness and patterns of asRNA transcription during cell growth and adaptation to different environments. In our work, we found that up to 93.5% of the annotated genes in the genome have sequencing reads mapped to their antisense strands; however, some of them may be transcriptional noise. To identify authentic asRNAs, we invoked a rather rigorous criterion to call asRNAs and found that varying annotated genes had asRNA transcription depending on sampling time points and culture conditions. We predicted a total of 660 asRNAs expressed under different time points under five culture conditions. Compared to other studies (10–12, 26), our number of predicted asRNAs was relatively small, but our threshold of calling “what is an asRNA?” was higher than most studies did. To be qualified as predicted asRNAs, they had to be observed in at least two biological replicates from the same time point, have at least 5 raw reads each, and have a TSS signal available by either dRNA-seq or manual review. So, the 660 asRNAs might be a lower bound of the authentic asRNAs in the species, as it is highly unlikely that the same transcriptional noise can repeat itself exactly using our stringent standard.

We observed that the asRNA expression showed a time point- and culture condition-dependent manner, evidenced by that most of these asRNAs were only expressed in certain time points and culture conditions. The results indicated that antisense transcription in *E. coli* K12 is highly variable and dynamic depending on growth phases and culture conditions. All our predicted asRNAs were observed in at least two biological replicates at the same time point, and many of them were all transcribed at certain time points under different culture conditions. Our findings of the high dependency of the transcription of the predicted asRNAs on the environmental changes, strongly indicate that these asRNAs may have important biological functions for the bacterium to adapt to different conditions.

We classified the transcriptional events of a gene locus in six possible modes according to its relative asRNA level to the mRNA level. The proportions of genes in these modes were highly variable, depending on the growth phases and culture conditions, and many gene loci changed their transcriptional modes at different time points under the culture conditions. It has been shown that asRNAs can either down- or up-regulate the expression of the cognate genes (20, 35) that are involved in important processes such as DNA replication, stress responses, and iron transport (20). Our results support these earlier observations as we found that for some genes such as *sulA* (Fig.6A), relatively elevated antisense transcription was accompanied by decreased expression of the protein, while for some other genes such as *uspF*, relatively elevated antisense transcription was concomitant with increased expression of the protein (Fig.6B). Therefore, the transcriptional modes that a gene locus adopts at different time points and culture conditions can be well explained by the known functions of the gene. All these results suggest that asRNAs may play a crucial role during the bacterium’s growth and adaptation to environmental changes.

Several mechanisms have been proposed to explain how asRNA can affect gene expression, including transcriptional interference, alteration of mRNA stability, and modulation of translation (6). Although the detailed molecular mechanisms remain to be elucidated, our results are in excellent agreement with the threshold linear response model for the stoichiometric interaction between asRNAs and cognate mRNAs (43, 44). According to this model, the formation of a duplex between an mRNA and its cognate asRNA will either decrease or increase the transcription, translation, or stability of the mRNA. Therefore, an increase in the relative level of an asRNA disrupts an otherwise strong correlation between the protein and mRNA levels. This model also is consistent with our finding that the lengths of asRNAs are irrelevant to their functions because only a short asRNA is needed to exert its function regardless of the length of the gene according to the threshold linear response model.

## MATERIALS & METHODS

### Bacterial culture and collection

A stock of *E. coli* K12 MG1655 strain stored in glycerol buffer at -80C⁰ was thawed at room temperature. 25µl of the *E. coli* stock was mixed with 2.5ml LB in a 10ml culture tube and incubated overnight at 37 C⁰ and 125rpm. On the second day, 2.5 ml *E. coli* culture was added to 250 ml fresh LB medium and cultured at 37 C⁰ and 125rpm. For the LB culture condition, cells were collected when an optical density at 600 nm (OD_600_) reached 0.5, 1.0, 20., 3.0, and 4.0. For the other four culture conditions, when OD_600_ of the culture in LB reached 1.0, cells were collected by spinning at 3500g for 20 minutes at room temperature, and then suspended and transferred to the desired mediums with the same volume (150ml). The other four stress culture medium were formulated as follows: MOPS (100ml of 10X MOPS mixture, 880ml of sterile H2O, 10ml (0.132M) KH2PO4, and 10ml of 20% glucose, Teknova, Hollister, CA, USA), HS (MOPS incubated at 48°C and 250 rpm), MP (MOPS without KH_2_PO_4_) for phosphorus starvation, and MC (MOPS without glucose) for glucose starvation.

At the designed time points for each culture condition, 10 ml *E. coli* culture was taken and added to a centrifuge tube containing 5 ml cold RNAlater (Thermo Fisher Scientific, Waltham, MA, USA). After centrifuging for 5000g at 4 C⁰ for 5 minutes, we discarded the supernatant and added 1 ml cold RNAlater. The cells were allocated to vials after re-suspension and precipitation and stored at -80C⁰ until use. mRNA isolation and enrichment were performed using MICROBExpress™ Kit (Thermo Fisher Scientific, Waltham, MA, USA). In general, the 16S and 23S rRNAs were captured by Oligonucleotide Mix and MagBeads in the MICROBExpress™ Kit and mRNAs were recovered by Ethanol precipitation as previously described (13).

### Extraction and enrichment of mRNAs

Total RNA was isolated using mirVana™ miRNA Isolation Kit (Thermo Fisher Scientific, Waltham, MA, USA). Briefly, after using lysozyme to release the cell content and a homogenizer to physically further break cells, we followed the vendor’s instruction to extract and purify total RNAs with the mirVana™ miRNA Isolation Kit. A round of DNase I treatment using RiboPureTM - Bacteria AM1925 (Thermo Fisher Scientific, Waltham, MA, USA) was applied to remove DNA residual from the experiment. Next, for the dRNA-seq procedure, we split the RNAs into two equal halves. We added Terminator™ 5-Phosphate-Dependent Exonuclease (Lucigen, Wisconsin, USA) into one part for the +TEX sample following the user’s manual and skipped this step for the other half for -TEX sample. Cleanup was done using RNeasy MinElute Cleanup kit 50x (Qiagen, Hilden, Germany).

### Construction of dRNA-seq libraries

The dRNA-seq libraries were constructed using Illumina Small RNA Sample Prep Kit (Illumina, San Diego, CA, USA) following the vendor’s instruction with some modifications. Briefly, after the purified mRNA was fragmented using an RNA fragmentation kit (Ambion, Austin, TX, USA), the fragmented RNA was treated with Antarctic phosphatase (NEB) to remove the 5’-tri-phosphate groups of RNAs with an intact 5’-end. A mono-phosphate group was then added back to the 5’-end of RNAs by polynucleotide kinase (PNK, NEB) in the presence of 10mM ATP. The sRNA 3’ Adaptor (5’/5rApp/ ATCTCGTATGCCGTCTTCTGCTTG /3ddC/) was ligated to the 3’-end of fragmented RNAs using truncated T4 ligase 2 (NEB), and the SRnA 5’ RNA adaptor (5’GUUCAGAGUUCUACAGUCCGACGAUC) was ligated to the 5’-end of fragmented RNAs using T4 ligase. Two rounds of PCR application were performed, and samples were cleaned up by AMPure XP before ready for sequencing. For our experiments, sequencing was done on an Illumina HiSeq 2000 platform (Illumina, San Diego, CA, USA) at David H. Murdock Research Institute of the North Carolina Research Campus (Kannapolis, NC, USA), and we constructed the directional RNA-seq libraries using Illumina’s TruSeq Small RNA Sample Prep Kit (Illumina, San Diego, CA, USA), so that multiplex sequencing can be achieved by using the barcoded PCR primers.

### Identification of asRNA

We trimmed adapter sequences using Cutadapt (48) and assessed sequencing reads quality using FastQC (https://www.bioinformatics.babraham.ac.uk/projects/fastqc/). The adapter-trimmed and high quality reads were mapped to the *E. coli* K12 MG1655 genome using Bowtie2 (45). The number of sense/antisense reads mapped for each gene was calculated by the program featureCounts (49). To identify asRNA, we first converted the uniquely mapped reads to the graph file format (reads per nucleotide), which is similar to the .bigWig format used in the genome browser. We normalized these reads using a method analogous to total counts (42, 46). We pooled the biological replicates at the same time point together as they were highly correlated (Table S3), but the sample information was still used later for asRNA identification.

We then scanned the graph files and output the longest transcripts from both strands. To be qualified as an “authentic” transcript, it must be seen in at least two replicates at the same time point and there are at least 5 raw reads along the transcript in each replicate. By comparing the relative location of the transcripts to the gene annotation and incorporating the strand information, we can classify the transcripts as “sense transcripts”, “antisense transcripts”, “intergenic transcripts”, and “unidentifiable transcripts” referring to those in the gene overlapping regions thus they cannot be labeled as “sense” or “antisense”. We grouped the transcripts with the “antisense” label associated with the same gene/operon. By using the TSS information output by TSSPredator (34), we predicted a collection of antisense transcripts. We lastly had a round of manual review and those that passed the human check were reported as final asRNAs that were used for analysis in this work.

### UPLC and Tandem Mass Spectrometry Analysis

Fifty µg of total protein from each sample were separated on 10% Bis-Tris NuPAGE gels (Invitrogen, Carlsbad, CA, USA) with 6X sample buffer containing 300mM Tris-HCl, 0.01% (w/v) bromophenol blue, 15% (v/v) glycerol, 6% (w/v) SDS and 1% (v/v) β-mercaptoethanol after denaturation at 95°C for 5 minutes. Gels were stained with the GelCode® Blue stain reagent (Thermo Scientific, Rockford, IL, USA) after fixation using 50% methanol (v/v) with 7% acetic acid (v/v) for 5 min. After destaining with water, each gel lane was excised into twenty slices which were put into in-gel tryptic digestion and peptide extraction according to the method reported previously (47). The dried residues were resuspended in 25µL of 10% ACN (v/v) with 3% formic acid (v/v) for LC-MS/MS analysis. The LC-MS/MS system used consisted of an LTQ/Orbitrap-XL mass spectrometer (Thermo Scientific, Rockford, IL, USA) equipped with a Nanoacquity UPLC system (Waters, Milford, MA, USA). Peptides were separated on a reversed phase analytical column (Nanoacquity BEH C18, 1.7µm, 150mm, Waters, Milford, MA, USA) combined with a trap column (Nanoacquity, Waters, Milford, MA, USA). Good chromatographic separation was observed with an 80 min linear gradient consisting of mobile phases solvent A (0.1% formic acid in water) and solvent B (0.1% formic acid in ACN) where the gradient was from 5% B at 0 min to 40% B at 65 min at 0.35µL/min of flow rate. MS spectra were acquired by data dependent scans consisting of MS/MS scans of the eight most intense ions from the full MS scan with dynamic exclusion of 30 seconds.

### Immunoblotting analysis

Twenty µg protein from each sample were separated by 8% sodium dodecyl sulfate (SDS)-polyacrylamide gel (SDS-PAGE) and transferred to nitrocellulose membranes. The membrane was blocked in 7% fat-free dry milk in TBS containing 0.2% Tween-20 and probed with antibodies against interest proteins. The antibodies used in this study included rabbit anti-*Escherichia coli* antibodies for UspF (Mybiosource, San Diego, CA, USA), SulA (Mybiosource, San Diego, CA, USA) and GAPDH (abcam, Cambridge, MA, USA). The GAPDH protein is known to be stably expressed under different conditions (48) and was used as the control for the western blot experiments. These primary antibodies were applied to the membranes in a dilution of 1:2000. Following extensive washing, the membranes were incubated with an HRP-labeled secondary antibody. The blots were then developed using the ECL western blotting substrate (Thermo Scientific, Rockford, IL, USA). The levels of protein expression were semi-quantified by optical densitometry using Image J Software version 1.46 (49). The ratio between the net intensity of each sample to that of the GAPDH internal control was calculated and served as an index of relative expression of an interested protein.

### Data visualization

Data visualization was prepared with R (50) and R packages ggplot2 (51) and VennDiagram (52).

### Data accession number

All the RNA-seq datasets supporting the conclusions of this article are available in the Gene Expression Omnibus (GEO) repository, with accession numbers GSE48151 and GSE64021. All the mass spectrometry data are deposited to the ProteomeXchange Consortium via the PRIDE partner repository with the dataset identifier PXD001640 and DOI 10.6019/PXD001640.

## ACKNOWLEDGEMENTS

We would like to thank Dr. Jingyun Lee and Dr. Sunil Hwang for their assistance in UPLC and Tandem Mass Spectrometry Analysis and Harry Lee for his contribution in preliminary data analysis.

