## Supplementary material for "Antisense transcription and its roles in adaption to environmental stress in *E. coli*": Figure S1

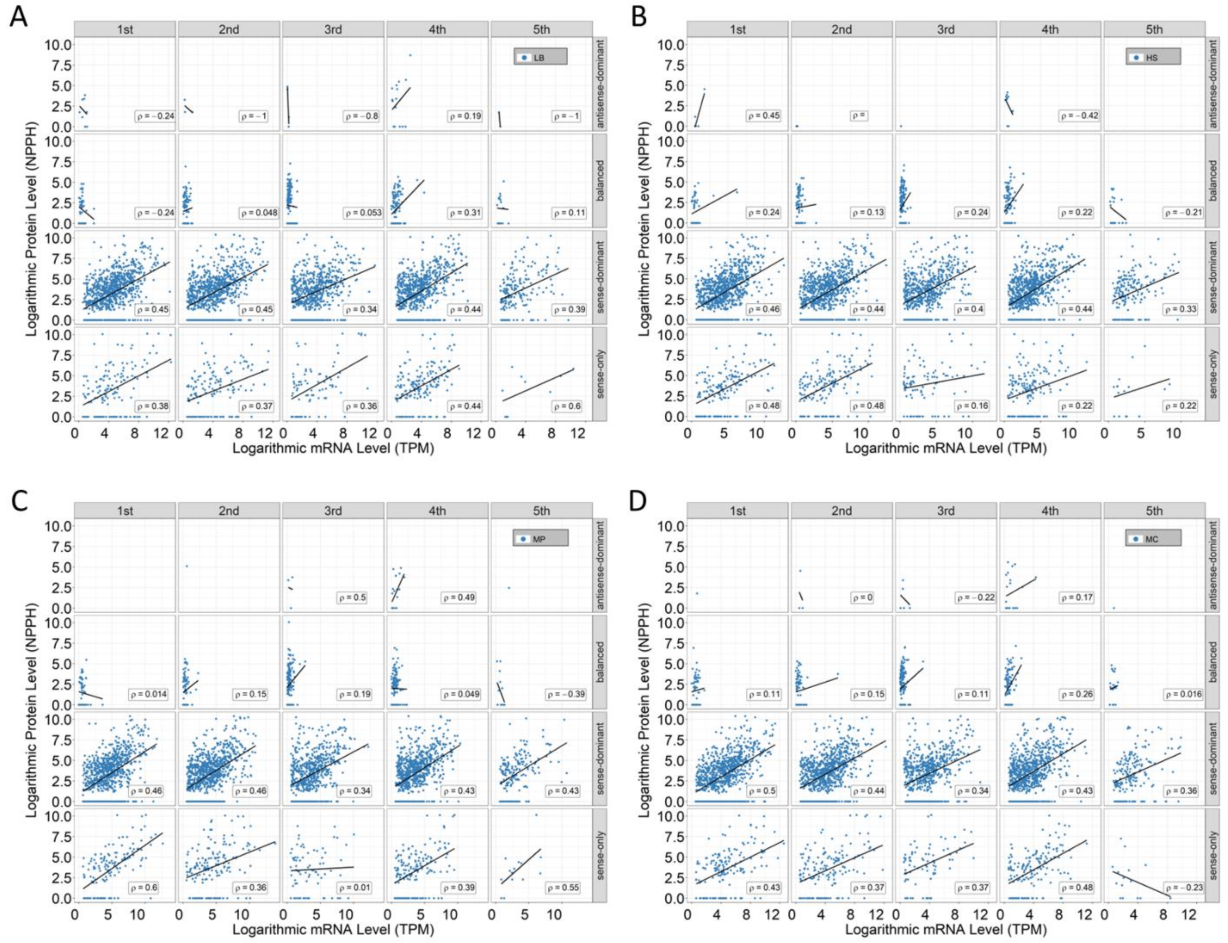


**Figure S1 Relationship between the sense transcription levels and protein levels of genes under four culture conditions.** Correlations between protein levels and mRNA levels of genes in different transcriptional modes (rows) and at different time points (columns) under (A) LB, (B) HS, (C) MP, and (D) MC conditions. Each data point represents one gene. The Spearman correlation coefficients ρ between the protein levels and mRNA levels of genes are shown in each plot.
